# Defining the impact of deep brain stimulation contact size and shape on neural selectivity

**DOI:** 10.1101/2020.06.11.147157

**Authors:** Daria Nesterovich Anderson, Alan D. Dorval, John D. Rolston, Stefan M. Pulst, Collin J Anderson

## Abstract

**Background:** Understanding neural selectivity is essential for optimizing medical applications of deep brain stimulation (DBS). We previously showed that modulation of the DBS waveform can induce changes in orientation-based selectivity, and that lengthening of DBS pulses or directional segmentation can reduce preferential selectivity for large axons. In this work, we sought to answer a simple, but important question: how do the size and shape of the contact influence neural selectivity?

**Methods:** We created multicompartment neuron models for several axon diameters and used finite element modeling with standard-sized cylindrical leads to determine the effects on changing contact size and shape on axon activation profiles and volumes of tissue activated. Contacts ranged in size from 0.04 to 16 mm^2^, compared with a standard size of 6 mm^2^.

**Results:** We found that changes in contact size induce substantial changes in orientation-based selectivity in the context of a cylindrical lead, and rectangular shaping of the contact can alter this selectivity. Smaller contact sizes were more effective in constraining neural activation to small, nearby axons representative of grey matter. However, micro-scale contacts enable only limited spread of neural activation before exceeding standard charge density limitations; further, energetic efficiency is optimized by somewhat larger contacts.

**Interpretations:** Small-scale contacts are optimal for constraining stimulation in nearby grey matter and avoiding orientation-selective activation. However, given charge density limitations and energy inefficiency of micro-scale contacts, our results suggest that contacts about half the size of those on segmented clinical leads may optimize efficiency and charge density limitation avoidance.

## Introduction

Recently, the deep brain stimulation (DBS) field has advanced in a number of directions with the goal of maximizing therapeutic benefit while minimizing side effects. Numerous reports have been published describing the modulation of DBS pulse width to induce changes in axon size selectivity^1–4^. At the same time, advancements have been made in DBS technology and programming. Segmented electrodes have been proposed and built, enabling the directional steering of stimulation to varying degrees of precision^2,5–7^, and optimization algorithms have been created and validated^8–11^, often with the goal of avoidance of maximizing stimulation within a specified target, such as the subthalamic nucleus, or avoidance of other targets, such as the internal capsule. Further, several groups have investigated modifications in charge balancing and leading pulse polarity to optimize neural selectivity^12,13^. Finally, several groups have investigated methods of generating orientation-specific selectivity, whether through the use of multiple electrodes^14,15^, multiple contact cathode configurations^16^, or alterations in the charge balancing pulse or leading pulse polarity^12^.

Many reports on the modulation of DBS axon size-specific selectivity have focused on changing the pulse width to achieve desired effects in selectivity. However, we recently reported that using directionally segmented contacts may be a more optimal means of focusing the effect of deep brain stimulation on nearby small axons, reducing the inherent preferential selectivity for large axons, which has been a goal of numerous groups in improving side effect avoidance^1^. Further, we recently proposed a novel, multiresolution DBS electrode design with the potential for contacts to vary in size and shape^2^, and the simple modeling studies reported therein provided further evidence that modulation of contact size may be exploitable in order to modulate neural selectivity. Based on our previous work and the importance of neural selectivity, we sought to more precisely define the impact of DBS contact size and shape on fiber size and orientation selectivity.

In this reported work, we modeled the segmentation of a standard-sized cylindrical DBS electrode into a total of 400 contacts, each 200 × 200 μm in size. In so doing, we were able to precisely examine the effects of contact size and shape on several aspects of selectivity, particularly orientation and fiber size selectivity. We expand the relationship of contact size to axon-size selectivity from our previous work, define how contact size can dramatically alter orientation-based selectivity for different orientations of passing fibers, show how to exploit rectangular shapes to increase orientation selectivity, and define the relationship between maximum spread and contact size, constrained by charge density limitations.

## Methods

### Overview

We modeled the influence of monopolar, biphasic cathode-leading extracellular stimulation on neuronal activation using multicompartment NEURON models, with fibers arranged vertically and tangentially (horizontally) around the electrode. We determined activation profiles for 2.0, 5.7, and 10 μm diameter axons using a theoretical cylindrical DBS electrode design consisting of 400 contacts arranged as 20 contacts vertically by 20 contacts horizontally wrapping around the lead, each contact 200 × 200 μm in size (Fig. 1). Given a circumference of 4 mm, this resulted in a 1.27 mm diameter lead, identical to standard cylindrical DBS leads. We modeled in a current-controlled format for easy computation of impedance, ranging amplitudes from 0 to 20 mA, with a fixed pulse width of 60 μs. We analyzed the effect of contact size and shape on axon size-specific and orientation-specific selectivity by comparing activation profiles of differently oriented electrodes of all three modeled fiber sizes.

**Figure 1:**
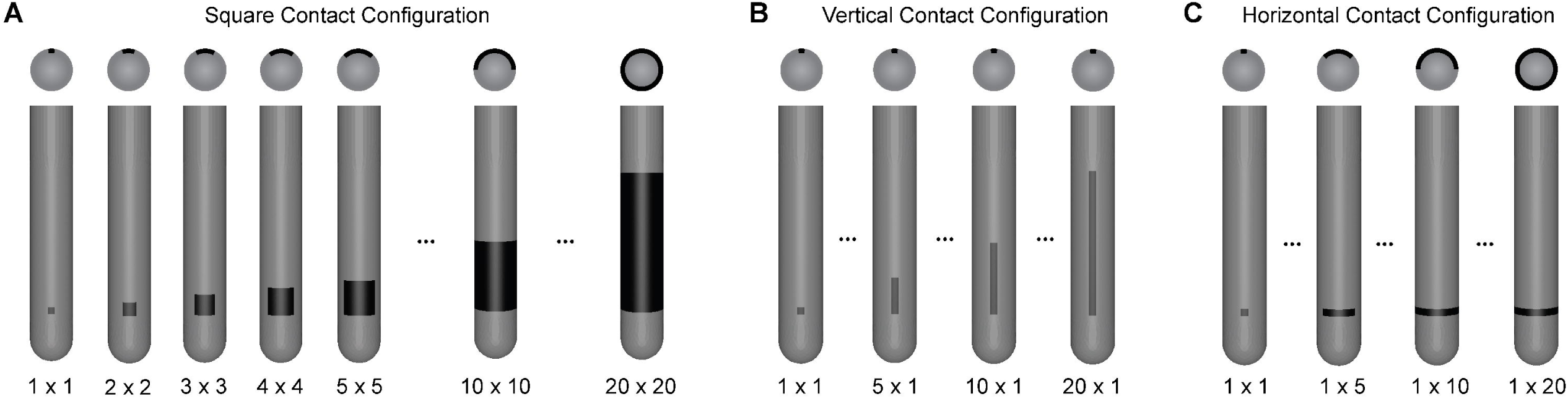
Electrode model enabling micro-scale modifications in contact size and shape. We modeled cylindrical leads with a standard 1.27 mm diameter overlaid with a 20 × 20 grid of square micro-contacts, each 200 µm x 200 µm. For each simulation, axons were positioned perpendicularly out from the center of the active contact. **A**. We modeled “square” contacts wrapping around the face of the lead from 1 × 1 to 20 × 20 contacts, thus 0.04 mm^2^ to 16 mm^2^ in surface area. These FEMs were used in figs. 2, 3, 5, and 6. The views shown are axial (top) and coronal (bottom). **B-C**. We modeled vertically (B) and horizontally (C) oriented rectangular contacts, ranging from 1 × 1 to 1 × 20 contacts, thus 0.04 mm^2^ to 0.8 mm^2^ in surface area. While horizontally oriented rectangular contacts wrap around the lead more with increased size, vertically oriented contacts do not. These FEMs were used in fig. 4. Note the comparability of our modeled contacts with clinical contacts, which are ∼4 mm in circumference and 1.5 mm in height.

### Finite Element Model

We modeled as previously described using the finite element method (FEM)^1,2^, implemented in SCIRun 5.0 (Scientific Computing and Imaging (SCI), Institute, University of Utah, Salt Lake City, UT), to solve the bioelectric field problem. We set electrode contacts as ideal conductors and electrode shafts as ideal insulators. We modeled an isotropic medium around the electrode, with 0.1 S/m conductivity^17^. Notably, we repeated experiments with different conductivities, including and excluding an encapsulation layer and found that while activation thresholds changed in response to changes in conductivity layer, the conclusions were similar regardless of conditions. We implemented a Dirichlet boundary condition at the outer edge of the model to serve as a distant ground ^1,8,18^ (100 mm x 100 mm x 100 mm) and solved for the electric potential solution.

### Neuron Modeling

We quantified neuronal response to extracellular stimulation through multicompartment axon modeling in the Neuron 7.7 environment. We used the McIntyre Richardson Grill (MRG) model^19^ for 5.7 μm diameter myelinated axons and modified parameters of this model for 2.0 μm and 10 μm myelinated axons^1,2,20,21^. We distributed tangential and vertical neurons evenly around the electrode in 0.2 mm increments from 0 to 10 mm in all directions. We interpolated electric potential solutions onto each node, paranode, and internode segment. We performed a binary search algorithm with cubic interpolation to determine the firing threshold within ∼0.001 mA resolution. We have previously reported that cathodic stimulation preferentially activates passing – i.e. vertical or horizontal – axons, while anodic stimulation preferentially activates orthogonal (radial) axons.^12^ Given our desire to examine size and shape under standard cathodic stimulation, we simulated the two primary directions of passing neurons, vertical and horizontal – longitudinal and latitudinal in spherical coordinates.

### Energy and Charge Density Calculations, Maximum Spread Modeling

We modeled under current controlled scenarios, and converted to voltage using FEM-derived impedances and Ohm’s Law, V = I x Z, where V is voltage, I is current, and Z is impedance. Energy is defined as V^2^ x F x PW x 1 sec / Z, where F is frequency and PW is pulse width. As contact size is modulated, frequency of stimulation and pulse width remain constant, while the impedance is derived from the FEM solution based on contact size and the constant tissue conductivity. Charge density was computed as (V x PW) / (Z x SA), where SA is defined as surface area. When determining maximum spread from the electrode given charge density limitation, we solved for the voltage resulting in a charge density of 30 μC/cm^2^ and determined the maximum distance at which an axon of each diameter at each orientation could be activated by said voltage.

### Quantifying Axon Size- and Orientation-Specific Selectivity

We quantified the effects of changes in contact size and shape on neural selectivity in several ways. We began by finding the maximum distances for which 2.0, 5.7, and 10.0 μm axons in both vertical and horizontal orientations are activated by 60 μs, -1.0 mA pulses from a 200 μm x 200 μm contact. Maintaining 2.0, 5.7, and 10 μm axons at these specific distances from the electrode, we found activation thresholds for all three axons given varied contact sizes and shapes. Importantly, while the distance from the electrode was maintained, the modeled axons were always centered perpendicularly out from the center of the active contact. Given equivalent thresholds across axons with a small, square contact, we calculated ratios comparing activation thresholds of different axon sizes given larger square and rectangular contacts for both orientations. Further, we calculated ratios comparing activation thresholds for the differing orientations as contact size was increased and shape was changed.

### Volume of Tissue Activated

We generated volumes based on previous methodology^8,12,18^, in which we isosurfaced the maximum eigenvalue of the Hessian matrix of second spatial derivatives of the electric potential based on second difference vs. stimulation threshold fit functions. For each fiber diameter, we generated the fit functions using a two-point Power law based on the firing threshold and maximum second derivative for tangential neurons spaced 0.2 mm apart along an orthogonal line radiating from the contact center up to 10 mm away. We solved for the Hessian matrix over a 20 × 20 × 20 mm grid with 0.1 mm spacing.

## Results

### Contact size influences orientation-specific selectivity

We modeled changes in “square” contact sizes on a cylindrical electrode shaft from 200 to 4000 μm per side (Fig. 1A) and found activation thresholds in current and voltage for axons situated 2 mm from the center of the lead. We note that “square” refers to an identical width and height for a given contact, though the contact wraps around the cylindrical lead. Increases in contact size increased the amplitude threshold in terms of current for both horizontal (Fig. 2A) and vertical (Fig. 2B) axons, and thresholds increased more substantially with smaller axon sizes. Notably, larger diameter axons featured smaller activation thresholds at all contact sizes, as expected. While current thresholds always increased for all axon sizes with increased contact surface area, voltage threshold relationships were more complex. For both horizontal (Fig. 2C) and vertical (Fig. 2D) axons, increasing from very small contacts (0.04 mm^2^ minimum) leads to an initial decrease in voltage threshold for all axon sizes at both orientations. Horizontal axons of all sizes quickly stabilized in threshold amplitude, remaining nearly unchanged with increased contact size past ∼3 mm^2^. However, activation thresholds for vertical axons increased in voltage past ∼3 mm^2^, particularly for 2.0 µm axons. Notably, substantial decreases in impedance as contact sizes increased led to the differing relationships between current and voltage thresholds with respect to contact size (see section on impedance below). Taking ratios of vertical / horizontal axon activation thresholds (Fig. 2E), we find that increase in contact size monotonically increased preference for horizontal fibers.

**Figure 2:**
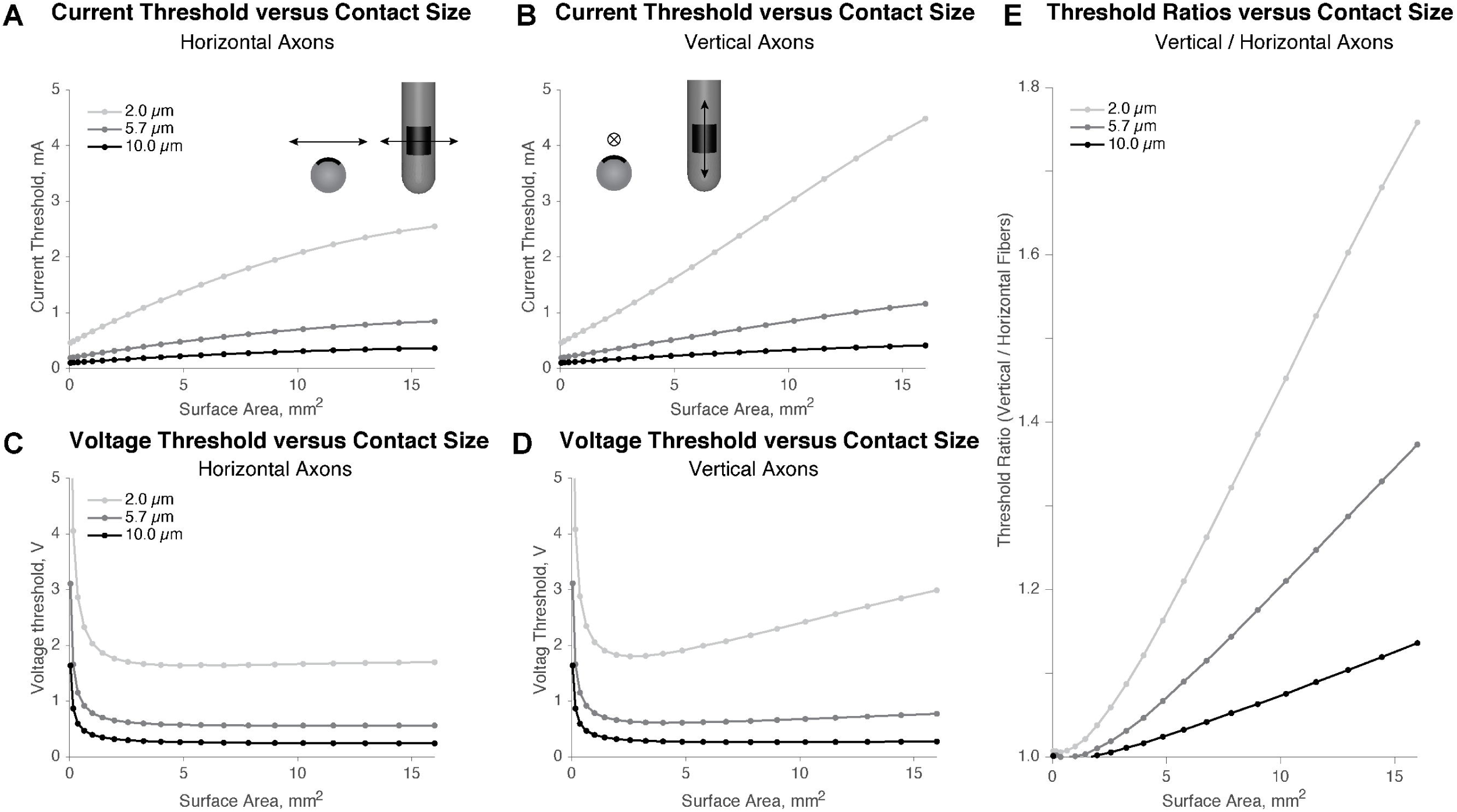
Contact size influences orientation-specific neural selectivity on a cylindrical lead. **A-B**. As the surface area of the “square” electrode contact increases, the current required to activate axons 2 mm from the electrode center of each size increases. Axons oriented horizontally (A) increase in threshold less rapidly than those oriented vertically (B) with increased contact size, particularly those with smaller diameter. **C-D**. Given much greater impedance for smaller contacts (see Fig. 6), the required voltage to activate axons of both orientations 2 mm from the electrode center initially decreases with increased size. However, while horizontal axons (C) plateau in voltage thresholds with increased contact size, vertical axons (D) of moderate size increase slightly in voltage threshold with increased contact size, while small axons increase substantially. **E**. As contact surface area increases, the ratio of amplitude thresholds for neuronal activation increases substantially, particularly in small axons. Note that this ratio is identical for either current or voltage.

### Contact size influences fiber size-specific selectivity

We completed two experiments to determine the effects of contact size on fiber size-specific selectivity. First, we found the threshold amplitudes in terms of current and voltage for horizontally and vertically oriented neurons at 2 mm distance from the center of the lead. As surface area of “square” contacts wrapping around the lead increased, the preferential selectivity for large fibers increased for both horizontal (Fig. 3A) and vertical (Fig. 3B) fibers. However, the multiplicative differences of different size axon activation thresholds nearly stabilized at the point when the surface area reached that of standard contact sizes (6 mm^2^) for horizontal fibers, while it continued increasing for vertical fibers. Regardless, repeating this experiment at a variety of distances, we found a robust result that smaller contacts most optimally decreased preferential selectivity of larger fibers.

**Figure 3:**
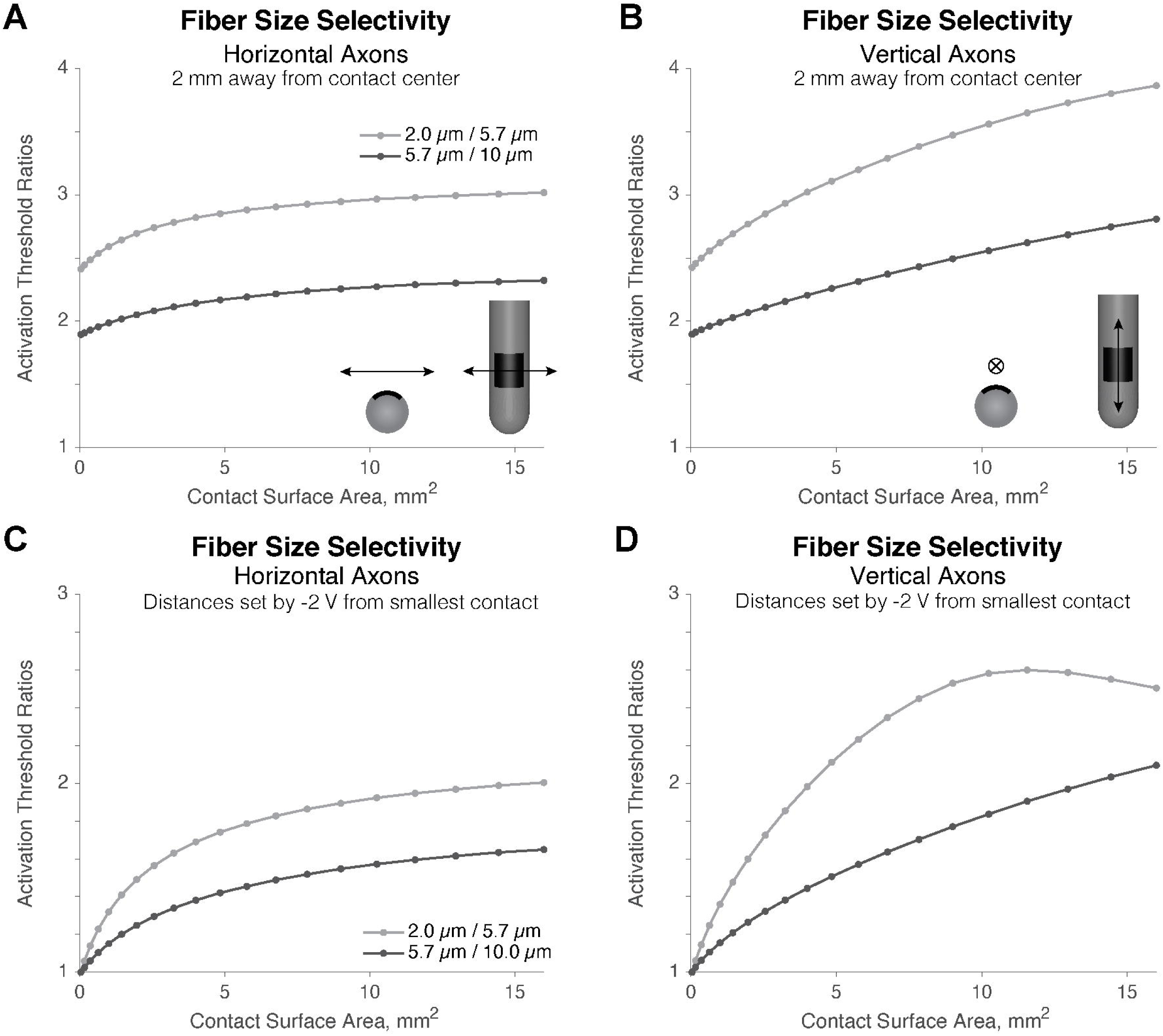
Smaller contact sizes reduce preferential selectivity for large fibers. **A-B**. We modeled axons of all sizes 2 mm from the center of the lead. For any contact size, smaller axons always had higher amplitude thresholds. However, for both horizontal (A) and vertical (B) axons, decreased contact size minimized the preferential selectivity for larger axons. Notably, while the ratios across thresholds nearly plateaued for horizontal fibers, they continued to increase substantially for vertical fibers **C-D**. We found the maximum distances where axons of each size were activated by -2.00 V, 60 µs pulses by a single 200 µm x 200 µm micro-contact, then found the minimum amplitudes to activate axons of all sizes at these given distances with increased contact sizes. For both horizontal (C) and vertical (D) fibers, increased contact sizes led to preferential selectivity for the larger, more distant fiber. Thus, in scenarios in which one wants to focus stimulation within nearby grey matter while avoiding distant white matter, smaller contacts are more effective. However, we note that, for vertical fibers, there is not a monotonic increase in preferential selectivity for the larger fiber with increased contact size.

Next, we sought to define the impact of contact size on the selectivity of nearby small axons versus distant, large axons. Thus, we found the maximum distances at which 2.0, 5.7, and 10.0 µm axons were activated by 60 µs, -2.00 V pulses from a single micro-contact (200 µm by 200 µm, i.e. a total surface area of 0.04 mm^2^). Vertical and horizontal axons of all sizes exhibited no preferential selectivity at such a small contact size – thus, the maximum distances were approximately 1.43 mm, 1.75 mm, and 2.18 mm from the electrode center for 2.0, 5.7 and 10.0 µm axons, respectively. At these axon locations, we found new activation thresholds for each axon size at each contact size. Horizontal fibers (Fig. 3C) showed minimal difference from the above relationship in terms of larger contacts preferentially selecting for large axons.

However, while vertical fibers (Fig. 3D) showed increased large fiber selectivity to a certain point with increased contact surface area, the trend eventually reversed, with very large contacts selecting for distant, 5.7 µm fibers over 2.0 µm to a reduced extent when compared to slightly smaller contacts. Notably, we repeated these simulations at numerous distances from the electrode, finding similar relationships at all.

### Contact shape influences fiber orientation selectivity

We modeled changes in contact size, using 200 μm wide contacts ranging from 200 to 4000 μm in height and 200 μm tall contacts ranging from 200 to 4000 μm in width (Fig 1. B,C). Horizontal axons were activated with reduced voltage thresholds from horizontal contacts of increased surface area until stabilizing at ∼1.5 mm in length (Fig. 4A), while they continued decreasing and reached lower thresholds in response to stimulation from vertically oriented contacts (Fig. 4B). Vertical axons stabilized at only slightly smaller thresholds than horizontal axons when stimulated by increasing sizes of horizontally oriented contacts (Fig. 4C). However, while 5.7 and 10.0 µm vertical axons stabilized in response to increasing length of vertically oriented contacts, 2.0 µm axons reached a minimum threshold with 1.8 mm tall contacts before continued lengthening led to increased thresholds. As contacts become more horizontally oriented, there is a slight preferential activation of vertical axons over horizontal axons (Fig. 4E). However, vertically oriented contacts lead to substantially greater preference for horizontal axons than horizontally oriented contacts do for vertical axons (Fig. 4F).

**Figure 4:**
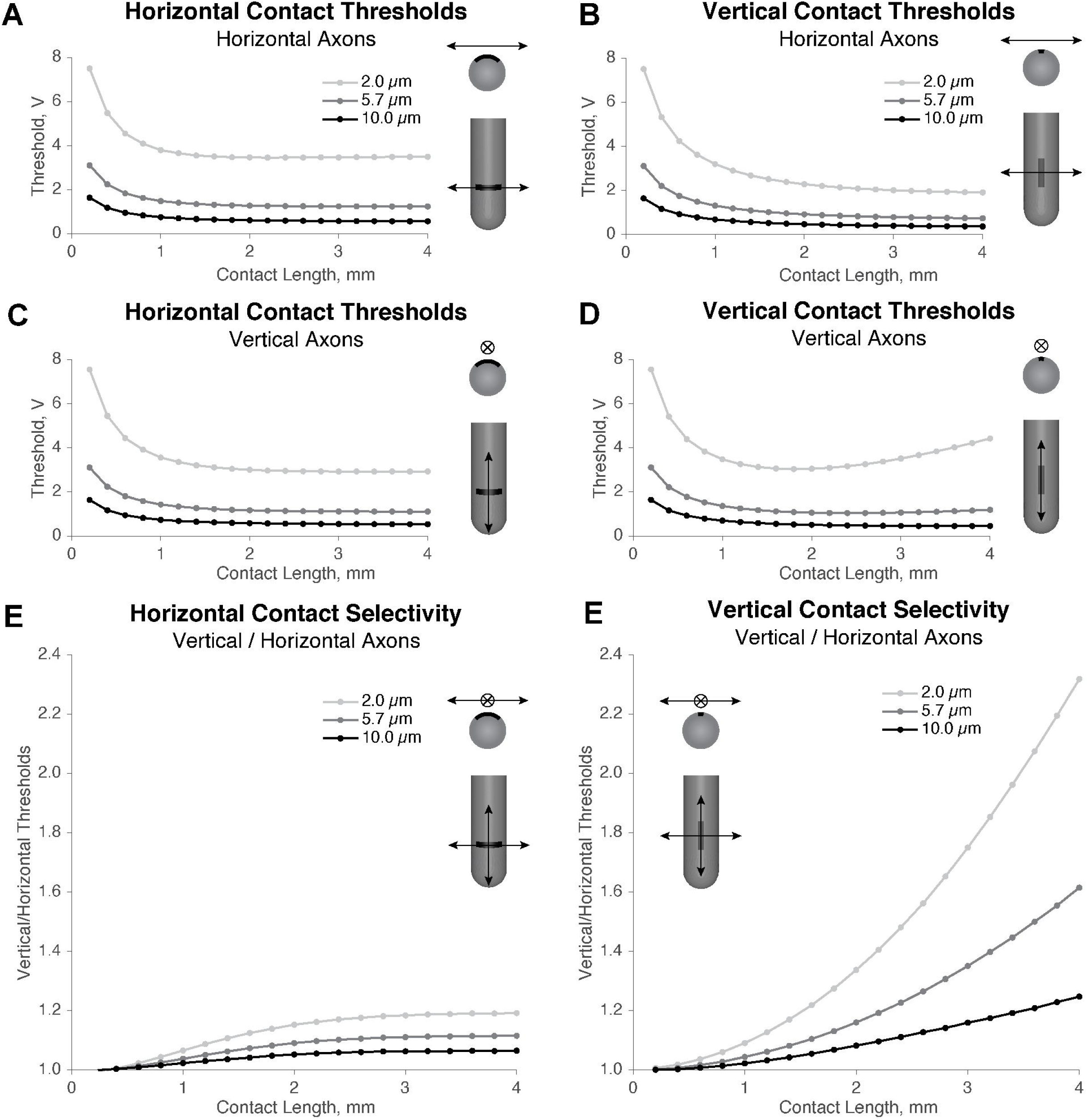
Contact shape influences orientation-specific selectivity on a cylindrical lead. **A**. Horizontally oriented rectangular contacts of increasing length decrease the threshold for horizontal axons of all sizes until stabilizing at ∼1.5 mm in length. **B**. Vertically oriented rectangular contacts decrease the threshold for horizontal axons beyond that seen in A. **C**. Horizontally oriented contacts of increasing length decrease the threshold for vertical axons until stabilizing at ∼1.5 mm in length. **D**. 5.7 and 10.0 µm vertical axons are stable in response to increasing vertical contact length past ∼1.5 mm length, but 2.0 µm axons, modeling grey matter axons, increase in threshold past ∼1.8 mm vertical contact length. **E**. Horizontally oriented contacts induce a small preferential selectivity for vertical neurons with increased size. **F**. Long, vertically oriented contacts induce substantial selectivity for horizontal axons, with increased orientation-specific selectivity with decreased axon size. We interpret that, since vertically oriented contacts do not wrap around the lead, they are better able to hold a long portion of vertical axons at fixed electric potential, decreasing the second spatial difference, which correlates with activation.

### Contact size influences the shape and volume of tissue activated

Changes in contact size led to substantial changes in the shaping of neuronal activation (Fig. 5A) under a current-controlled scenario, with 1 mA of current applied on all contact sizes. However, at least in the context of an isotropic medium, the volume of tissue activated was only minimally changed by changes in contact size (Fig. 5B), which supports the idea that changes in impedance — due to contact size, as explored in the next sub-section — do not substantially alter the VTA activated under a current-controlled scenario. That stated, normalizing to the contact size closest to a standard clinical contact (5.76 mm^2^) clearly shows a subtle relationship between the contact size and the volume of tissue activated (Fig. 5C). Notably, smaller than standard contact sizes, roughly on the order of clinical segmented contacts, optimize the selectivity for 2.0 µm fibers in the context of VTA, while increasing the contact sizes past standard clinical contacts increases selectivity for large fibers. Under a voltage-controlled scenario, increases in voltage increase the volume of tissue activated (Fig. 5D, E). Similar to the current-control scenario, normalizing to the contact size closest to a standard clinical contact (5.76 mm^2^ shows that smaller-than-standard contacts improve the selectivity for 2.0 µm fibers, while larger contacts increase preferential selectivity for larger fibers (Fig. 5F).

**Figure 5:**
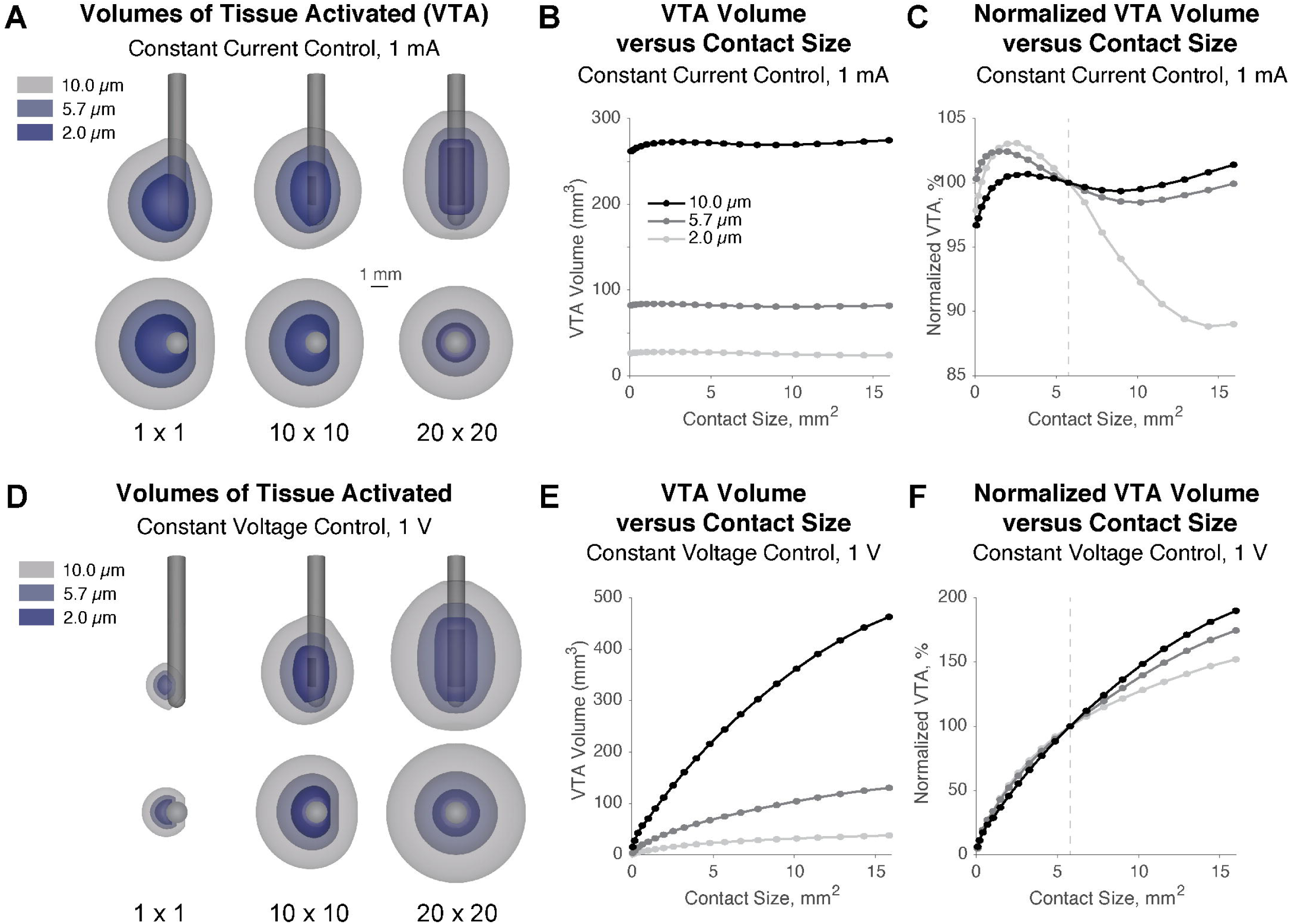
Contact size influences the volume of tissue activated. **A**. VTAs are shown under a constant-current scenario, 1.0 mA, for “square” contacts with surface areas of 0.04, 4.0, and 16.0 mm^2^. VTAs were generated from a sagittal (top) and axial (bottom) perspective for all 3 fiber sizes studied. **B**. Under constant current, changes in contact size yield only very minimal fluctuations in VTA. **C**. Normalizing the VTA to a nearly standard contact surface area (5.76 mm^2^) shows a subtle relationship between contact size and VTA. Notably, smaller-than-standard contacts decrease the preferential selectivity for larger VTAs in large fibers, while larger contacts increase the preferential selectivity for larger VTAs in larger fibers. **D**. VTAs are shown, as in A, but under a constant-voltage scenario. **E**. Increasing the contact size results in a larger VTA for all fiber sizes under constant-voltage conditions. **F**. Normalizing the VTA to a nearly standard contact surface area (5.76 mm^2^) shows that, with constant voltage, smaller-than-standard contacts are optimal for decreasing the preferential selectivity for large fibers, while large contacts minimize the spread over small fibers while preferentially selecting even larger VTAs for large fibers.

### Contact size limits the achievable spread of activation through charge density implications

Increased contact size led to decreased impedance (Fig. 6A) with a 5.76 mm^2^ contact exhibiting 1.08 kΩ impedance, consistent with standard cylindrical leads, and a 1.96 mm^2^ contact exhibiting 2.03 kΩ impedance, consistent with standard segmented leads.^22^ As the contact size decreases, the maximum acceptable amplitude, as limited by charge density, decreases. Here, we use the standard limitation of 30 µC/cm^2^ and find that the maximum possible current (and corresponding voltage) quickly exceeds the range of simulation parameters we tested, given we set a maximum stimulation amplitude of 20 mA (Fig. 6B). Given the relationship between contact size and the maximum current that can be applied before charge density limitations are met, we determined how contact size influences the maximum spread of activation achievable from the center of the electrode lead (Fig. 6C). With a single micro-scale contact of 0.04 mm^2^ surface area, stimulation can only reach 1.61 mm from the center of the lead (0.97 mm from the edge of the lead) in grey matter before charge density limitations are exceeded. Thus, micro-scale contacts may be quite ineffective at spreading stimulation much past an encapsulation layer. However, increasing to even a 1 mm^2^ contact enables activation 3.77 mm from the center of the lead in grey matter before reaching charge density limits. Notably, contact sizes in Fig. 6 are limited by the maximum distance from the electrode modeled (10 mm) and the maximum current modeled (20 mA).

**Figure 6:**
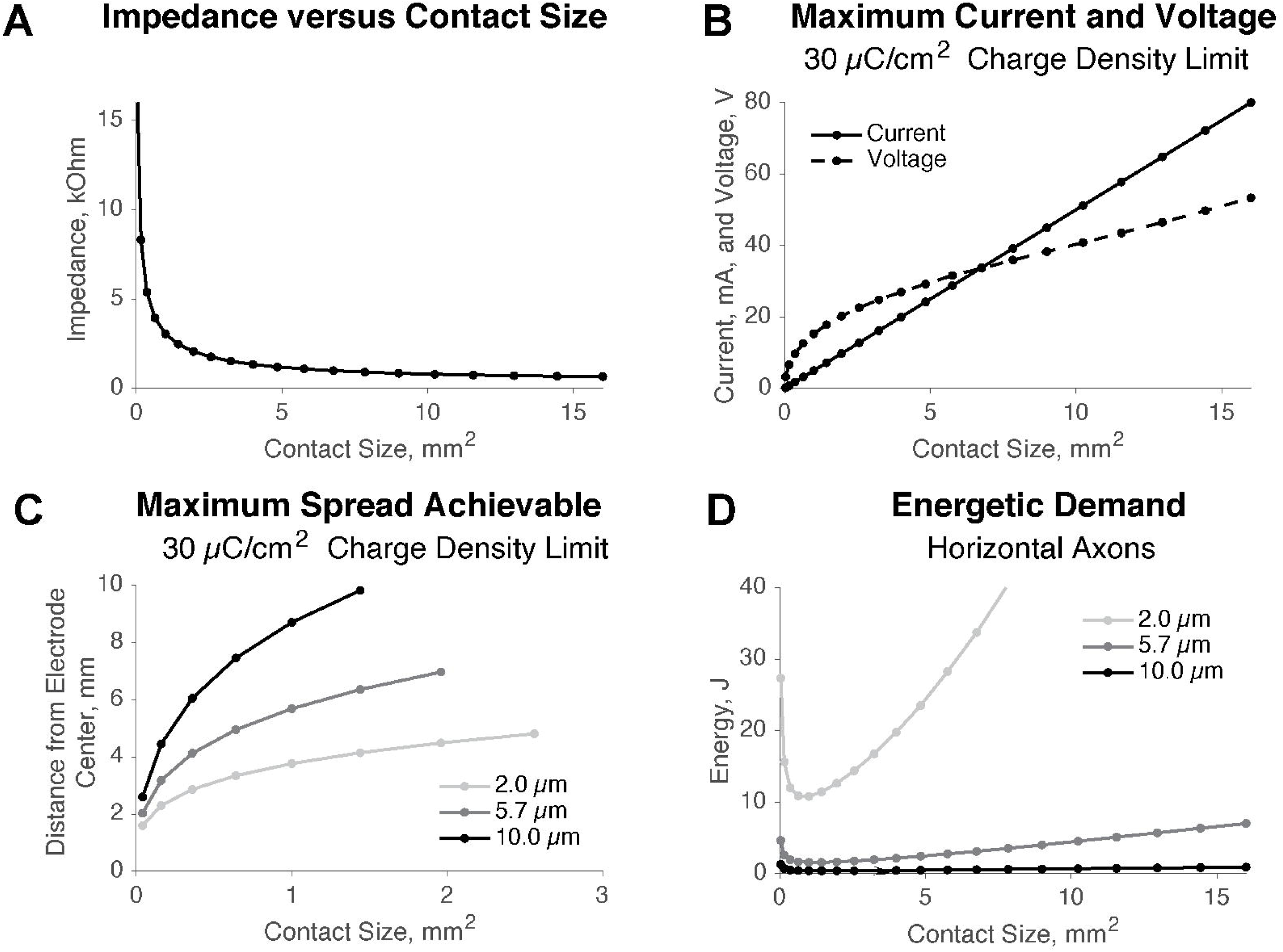
Micro-scale contacts are limited in utility by charge density and relative energetic inefficiency. **A**. Impedances decrease substantially as “square” contacts increases from micro-scale sizes to standard surface areas. **B**. Given a charge density limitation of 30 µC/cm^2^, contact surface area dictates the maximum current and voltage usable with 60 µs pulse width. Notably, with standard pulse widths, rather large amplitudes are possible. **C**. Given the amplitude limits determined in B., the maximum distance at which axons of various sizes can be activated is quite limited with micro-scale contacts. Notably, a single 200 µm x 200 µm micro-contact cannot activate grey matter size axons more than 1.61 mm from the center of the lead (0.97 mm from the edge of the lead) before charge density limits are exceeded. However, a ∼1 mm^2^ contact in an isotropic, medium-impedance model is predicted to enable grey matter activation 3.77 mm from the center of the lead. **D**. Energetic demand for grey matter stimulation is optimized by ∼1 mm^2^ of contact surface area.

### Relatively small – not micro-scale – contacts optimize energetic efficiency

Further hampering the utility of micro-scale contacts is the energetic inefficiency created by large voltage requirements on small contacts. With a goal of stimulating grey matter from a square contact, a surface area of ∼1 mm^2^ in size is predicted to minimize the energetic demand to stimulate an axon 2 mm from the center of the lead (Fig 5D). Further, we find that standard cylindrical contacts are, relatively speaking, fairly inefficient for grey matter stimulation, while large contacts are even more inefficient. Notably, energy calculations are derived from 130 Hz stimulation with 60 µs pulse widths for 1 second of total stimulation duration.

## Discussion

We and others have sought to define the impact of DBS pulse width, electrode configuration, and waveform on fiber size and orientation selectivity^1,3,12,23–25^. This work describes the impact of DBS contact size and shape on fiber size- and orientation-based selectivity. While standard electrodes preferentially select for larger fibers, we found that decreasing contact size from standard values could decrease this preference. Further, we found that non-square contacts can increase selectivity across passing axon orientations.

### Contact size modifies therapeutic window

We have previously argued that therapeutic window should be defined from a biophysical perspective rather than simply on amplitude, as modulating other DBS parameters, such as pulse width, can directly suppress or enhance the effects that amplitude has on the spread of neuronal activation. Indeed, when one wishes to focus the stimulation effect on small, nearby axons while avoiding stimulation of large, distant axons, the therapeutic window can be increased from a biophysical perspective despite a decreased window of possible amplitudes.^1^ Given a goal of increasing therapeutic window in this context, diminishing contact sizes to a certain point effectively increases the therapeutic window. However, grey matter is not always responsible for therapeutic benefit, so one must consider the target fiber when discussing therapeutic window. Regardless, the ability to use non-standard contact sizes could ultimately be useful in tuning neural selectivity towards specific fiber sizes in the future.

### Selective activation of horizontal or vertical axons, reduced or enhanced through modification of contact shape

With very small, square active contacts, passing fibers are selected approximately equally. However, it is notable that vertical axons are preferentially activated over horizontal axons with large, “square” contacts. As contacts grow very large, the horizontal component of the “square” contact wraps around the lead, resulting in less difference in the electric potential in the vertical plane versus the horizontal plane. The second difference of voltage serves as the driving force of neuronal activation; thus, we interpret that stimulating across the length of a vertically oriented axon with a large contact generates a more fixed voltage, decreasing the second difference. On the other hand, horizontally oriented axons will experience less stable voltages, generating a larger second differences, and thus, easier activation. Given the geometry of standard contacts, the preferential activation of vertical neurons by large “square” contacts seems unavoidable without making wider and wider leads, which would lead to more tissue damage.

The preferential selection of horizontal axons is enhanced by creating large, vertically-oriented rectangular contacts. We interpret that shaping the active contact vertically will reduce the driving force for activation in the parallel direction while not modifying the driving force in the perpendicular direction. Thus, the exploitation of orientation-specific selectivity through shaping of contacts demonstrates that our recently proposed multi-resolution electrodes could prove useful beyond what was discussed in our previous reports.^6,26^

### Decreasing contact size only increases efficiency up to a certain point; micro-scale contacts are not optimal

We previously showed that using 1 segmented contact on a standard directional lead is more energetically efficient than using 2 segmented contacts, which is, itself, slightly more efficient than using ring mode or a standard cylindrical contact. Thus, it may be expected that smaller contacts still may be even more efficient. Indeed, we found that contacts of ∼1/2 the surface area of a standard segmented contact on a directional electrode ought to be most energetically efficient, but decreasing past ∼1 mm^2^ in surface area leads to very large increases in voltage requirements. Given that energy depends on the square of the voltage, this makes contacts under ∼1 mm^2^ less efficient. If one has a goal of stimulating grey matter and avoiding distant white matter tracts that may be associated with side effects, it is clear that large contacts are not optimal. However, given energy inefficiency of micro-scale contacts and charge density limitations, it seems as though we may find benefit in the future from the flexibility of using micro-scale elements to create multi-scale contacts^2^, but using single micro-scale contacts would be ineffective. That stated, our work here leads to the prediction that having the ability to stimulate from active contacts with ∼1 mm^2^ surface area in the future may be beneficial.

### Charge density limits and their effect on the spread of neuronal activation

While smaller contacts are effective at decreasing the preferential selection of larger fibers, one must consider increased charge density. The typically recommended charge density limit of 30 μC/cm^2^ may be questionable^27^, but it remains certain that too high of charge density can damage tissue. Regardless of the chosen charge density limit, smaller contacts limit the overall volume of tissue activated possible before the specified limits are reached. Therefore, one must weigh neural selectivity versus charge density when considering contact sizes. Fortunately, smaller contacts have increased impedance, so the increase in charge density at a given voltage with a decrease in surface area is partially mitigated by impedance.

### Potential Caveats

It should be immediately noted that this manuscript was written with the intent of generalizability using simplified models that are non-specific to any individual patient. While this allows applicability to a number of scenarios of extracellular stimulation, specific numerical predictions outside of generalized relationships would need to be made with individualized patient models. For example, the precise numbers in the predictions of maximum spread vs. contact size are specific only to an isotropic model with a moderate impedance values. Thus, while it is reasonable to expect that the relationship will hold under a variety of implantation scenarios — and we have tested various conductivities and found qualitatively similar results — the specific results could not be used to make predictions as to the exact spread in a specific patient implant. Even in the context of a specific patient implant, one would expect these values to reduce over time if the conductivity decreases with additional encapsulation over time.

Further, axons are never oriented at precisely horizontal or vertical orientations to the electrode in reality, but have complex bends, which may initially be thought to complicate selectivity. However, as we have previously shown, selectivity results from simplified models, whether orientation-specific or fiber size-specific, have held up and can explain the observed activation patterns in patient-specific models.^1,12^

Additionally, when considering different methods of computing VTAs, we should note that there are numerous methods for computing a VTA. In this work, we focused exclusively on monopolar stimulation modeled with the axon model method. We have previously shown that all standard VTA methods for monopolar stimulation yield similar results^18^ — however, these similarities break down with bipolar stimulation. Thus, if one were to choose to expand our work to bipolar stimulation, it may also be necessary to test several methods of VTA computation.

All of the work here was completed in the context of a cylindrical DBS lead. Thus, the horizontal versus vertical orientation differences would not be expected to hold with other lead geometries unless the contact continues to wrap around the face of the lead. That said, implanting electrodes much wider than standard be highly impractical due to the tissue damage created by implantation, so it is likely that cylindrical leads or, at least, leads with similar cross-sectional area, will remain standard for the foreseeable future.

Finally, it should be noted that we only evaluated lead designs with one active contact, with 20 separate FEMs for each square contact design, and 38 additional FEMs for the remaining rectangular designs. Thus, modeling with other floating, conductive contacts present would result in slightly different results. However, we have previously evaluated a micro-scale contact design on a flat contact face with non-active contacts modeled as floating^2^, and in that scenario, there were no qualitative differences from modeling with active contacts. Thus, given the goal of generalizable results, it is unlikely that passive contacts would change the outcome.

## Conclusions

In this work, we use computational approaches to explicitly predict the effects of contact size and shape on several aspects of neuronal selectivity. We show that increased contact sizes may reduce activation threshold for a given axon to a certain point, but larger contacts may actually increase the threshold beyond that point. Contact size influences not only axon size-specific selectivity, but also orientation-specific selectivity when constrained within the context of cylindrical electrodes. We find that shaping of contacts can be exploited to enhance or reduce orientation-specific selectivity. We show that small contacts can be more energetically efficient, particularly when activation is desired in one specific direction, but this benefit is limited by a reduced maximum spread of activation with small contacts prior to running into charge density limitations. Thus, while modulations in contact size and shape will add additional dimensions to an already complex set of DBS programming variables, the future creation of multi-resolution clinical devices with more flexibility may enable substantial benefits in therapeutic outcome.

## Acknowledgements

This work was funded by NIH NINDS 2R37NS033123-14A1 (Pulst), NIH NINDS 1R21NS10479901 (Pulst), NIHNINDSK23NS114178(Rolston),NIHNINDSLRPUVWH4301(D.Anderson),NIHNINDS 1F32NS11432201 (D. Anderson), NSF-CAREER 1351112 (Dorval), Utah Neuroscience Initiative Collaborative Pilot Project Award (Pulst, Dorval), and National Ataxia Foundation Young Investigator Award (C. Anderson).

## Notes

### Competing Interest Statement

The authors have declared no competing interest.

